# Anti-Melanoma Differentiation-Associated protein 5 Auto-Antibodies Promote a Profibrotic Phenotype in a Human Lung Fibroblast Cell Line

**DOI:** 10.64898/2026.05.31.727600

**Authors:** Silvia Calandra, Maristella Maggi, Andrea Previtali, Luisa Iamele, Paola Cipriani, Luca Navarini, Roberto Giacomelli, Piero Ruscitti, Veronica Codullo, Giovanni Zanframundo, Claudia Scotti, Lorenzo Cavagna

## Abstract

Anti-melanoma differentiation-associated protein 5 (anti-MDA5) autoantibodies identify a distinct dermatomyositis subset frequently associated with rapidly progressive interstitial lung disease (RP-ILD). While these antibodies are established disease markers, their direct contribution to pulmonary fibrosis is poorly defined. This study investigated the pathogenic effects of patient-derived polyclonal anti-MDA5 antibodies on IMR-90 human lung fibroblasts. Recombinant human MDA5 protein was produced in HEK293F cells and utilized to selectively isolate autoantibodies from a patient’s plasma via affinity chromatography. Fibroblasts were stimulated with MDA5, anti-MDA5 antibodies, or both. Real-Time Cell Analysis (RTCA) showed a statistically significant increase in cell impedance following treatment with an MDA5–anti-MDA5 mixture compared with controls, accompanied by a reduction in cell doubling time. MTT assays showed that neither MDA5 nor anti-MDA5, nor their immunocomplex, exerted acute cytotoxic effects in cell culture. Direct cell counting revealed a significant increase in fibroblast proliferation in response to the MDA5–anti-MDA5 combination. Molecular characterization by RT-qPCR revealed a significant alteration of TLR2, TLR7, and endothelin-1 (ET-1) mRNA levels. ELISA assays detected an increased secretion of pro-collagen and type I interferons in culture supernatants. All these results were mainly, but not only, observed in the MDA5/anti-MDA5-exposed cells. Our results suggest that anti-MDA5 autoantibodies and MDA5 antigen complex are not merely disease biomarkers, but active pathogenic drivers that stimulate proliferation and pro-fibrotic responses in lung fibroblasts. This mechanism may contribute to the rapid tissue remodeling characteristic of RP-ILD, supporting the development of targeted therapeutic strategies to mitigate fibrosis in this high-mortality patient subset.

## 1. Introduction

Anti–melanoma differentiation-associated gene 5 (anti-MDA5) antibodies identify a distinct subgroup of dermatomyositis characterized by peculiar clinical and immunological features. Although initially described predominantly in Asian populations, anti-MDA5-associated disease is now increasingly recognized worldwide and represents a clinically relevant subset of dermatomyositis in both Asian and non-Asian patients^1,2^. Patients carrying these autoantibodies frequently develop clinically amyopathic or hypomyopathic dermatomyositis, often accompanied by constitutional symptoms, cutaneous ulcerations, and arthritis. However, the major determinant of prognosis and disease outcome in this subset is pulmonary involvement. Specifically, these patients are particularly prone to interstitial lung disease (ILD), especially rapidly progressive ILD (RP-ILD), which may develop within weeks to months from disease onset and is associated with high mortality despite aggressive immunosuppressive treatment ^3–6,1,2^.

The radiological patterns of pulmonary involvement in these patients are heterogeneous and most frequently include nonspecific interstitial pneumonia (NSIP) and organizing pneumonia (OP), either isolated or overlapping, although usual interstitial pneumonia (UIP)-like features have also been reported. Furthermore, anti-MDA5-positive patients frequently display a peculiar serological profile characterized by the absence of conventional autoimmune markers, such as the antinuclear antibodies, and by the positivity for anti-Ro52 antibodies in up to one-third of patients. The anti-Ro52 status had been associated with an increased risk of RP-ILD ^2^. In addition to pulmonary involvement, these patients often display systemic manifestations associated with chronic inflammation and tissue remodeling, suggesting that dysregulated immune responses may contribute to progressive fibrotic damage in multiple organs.

Melanoma differentiation-associated protein 5 (MDA5), encoded by the *IFIH1* gene, is a cytosolic RNA sensor belonging to the RIG-I-like receptor family and plays a crucial role in innate antiviral immunity. Upon recognition of long double-stranded viral RNA, MDA5 activates downstream signaling through mitochondrial antiviral-signaling protein (MAVS), promoting the activation of IRF3, IRF7, and NF-κB pathways. This leads to the production of type I interferons and pro-inflammatory cytokines ^7,8^. Persistent activation of these pathways has been associated with several autoimmune and interferon-mediated disorders, including anti-MDA5-associated dermatomyositis, where a strong interferon signature has been repeatedly described. Importantly, MDA5 is increasingly recognized not only as an antiviral sensor but also as a key player at the interface between innate immunity and autoinflammation. Aberrant activation of MDA5 signaling pathways has been implicated in the pathogenesis of autoimmune and inflammatory diseases, highlighting how dysregulated RNA sensing may contribute to chronic immune activation and tissue injury ^9^.

Despite the well-established diagnostic and prognostic relevance of anti-MDA5 antibodies, their direct contribution to disease pathogenesis remains poorly understood. Increasing evidence suggests that these autoantibodies are not simple biomarkers of immune dysregulation, but could actively participate in the inflammatory and fibrotic processes underlying tissue damage. Indeed, the clinical improvement observed after plasmapheresis in some cases of severe anti-MDA5-positive patients supports the hypothesis that circulating autoantibodies and immune complexes may exert direct pathogenic effects ^6^. Proposed mechanisms include the amplification of innate immune signaling, immune complex-mediated inflammation, endothelial dysfunction, and the activation of profibrotic pathways ^4,6^. Recent studies have further highlighted this complex landscape, suggesting that excessive activation of innate immune pathways, sustained interferon signaling, endothelial injury, and aberrant macrophage activation may all contribute to disease progression. Specifically, the inflammatory milieu associated with anti-MDA5 positivity may directly promote fibroproliferative responses within the lung microenvironment, ultimately contributing to the development and evolution of RP-ILD ^9,2^.

The marked association between anti-MDA5 autoantibodies, RP-ILD and lung fibrosis occurrence strongly suggests a pathogenic link between immune dysregulation and aberrant tissue remodeling. However, the molecular mechanisms underlying fibroblast activation and progression toward RP-ILD remain incompletely understood. The pathological process underlying ILD is characterized by persistent inflammation, activation of fibroblasts, excessive extracellular matrix deposition, and progressive remodeling of the pulmonary parenchyma, ultimately leading to compromised lung function. In this context, understanding whether anti-MDA5 antibodies directly contribute to fibroblast activation and fibrosis is of major interest. To investigate these potential profibrotic effects, the human lung fibroblast cell line IMR-90 was selected as the primary in vitro model. IMR-90 cells are normal human fetal lung fibroblasts widely used for studies of fibrosis and extracellular matrix remodeling due to their preserved fibroblastic phenotype and responsiveness to inflammatory and profibrotic stimuli ^10,11^.

The aim of the present study was therefore to investigate the potential profibrotic activity of patient-derived anti-MDA5 autoantibodies using this in vitro human lung fibroblast model. Specifically, we sought to evaluate whether purified polyclonal anti-MDA5 antibodies and MDA5–anti-MDA5 immune complexes could modulate fibroblast proliferation, viability, and the expression of genes involved in inflammation and extracellular matrix remodeling. By clarifying the biological effects of these autoantibodies, our study aims to better define their contribution to tissue fibrosis and pulmonary involvement in anti-MDA5-associated dermatomyositis, potentially providing insights to support the development of targeted therapeutic approaches.

## 2. Material and Methods

### Patient selection and ethical approval

A 48 old female patient with recent onset and treatment-naïve anti-MDA5 dermatomyositis, presenting with RP-ILD, admitted at the Rheumatology Unit of the IRCCS Policlinico S. Matteo of Pavia in 2025 was treated as per internal protocol with high dose corticosteroids (1 g of methylprednisolone in days 1,2, and 3 consecutive days, then 1 mg/kg/day with subsequent tapering, 3 plasma-exchanges (PLEX) on alternate days starting from day 4, followed by five days of High Dose Intravenous Immunoglobulins (2 mg/kg as a total dosage) starting from day 9, and then rituximab 1 g IV in days 15 and 30. Plasma obtained from the PLEX was immediately aliquoted and stored at −80°C until further processing. The study was approved by the local Ethical Committee of Pavia (P-201190088730; Prot. 20190094533).

### Cell Culture

The human fibroblast cell line IMR-90, derived from the lung tissue of a 16-week-old female fetus, was utilized for all in vitro experiments. Cells were cultured in Eagle’s Minimum Essential Medium, EMEM, supplemented with 10% fetal bovine serum (FBS), 1% penicillin/streptomycin, and 1% non-essential amino acids (NEAA). Cultures were maintained in a humidified atmosphere at 37°C with 5% CO_2_.

### MDA5 Protein Production and Purification

The HaloLink™ Resin (Promega) system was employed for the dual purpose of recombinant protein purification and subsequent autoantibody isolation. For protein purification, the Halo Link resin was incubated with the protein lysate obtained from HEK293F cells transfected with the encoding vector. Following a series of washes with HaloTag® Protein Purification Buffer (Phosphate Buffered Saline, pH 7.5, supplemented with 1 mM DTT and 0.005% IGEPAL® CA-630) to remove non-specifically bound cellular contaminants, MDA5 was cleaved using Halo-TEV protease (Promega; Madison, Wisconsin, United States) and eluted from the column. Its concentration was determined by microBCA (Sigma Aldrich; Saint Louis, United States) and utilized for direct in vitro treatment of IMR-90 cells or maintained linked to the resin to serve as an affinity matrix.

### Isolation of polyclonal antibodies

For autoantibody selection, the MDA5-functionalized resin was used to specifically capture anti-MDA5 polyclonal antibodies from the patient’s apheresis after processing it by centrifugation to remove cellular debris. Incubation was performed under constant rotation at 4°C overnight to ensure maximal binding. Following incubation, the resin-sample mixture was loaded onto a column (Amersham Biosciences, Little Chalfont Buckinghamshire, United Kingdom). The flow-through was collected by gravity, and the resin was subjected to extensive washing steps with HaloTag® Protein Purification Buffer to eliminate non-specific binders. The bound polyclonal antibodies were then eluted using a low-pH elution buffer (Glycine, pH 2.5-3.0). To preserve the structural integrity and biological activity of the purified antibodies, the eluate was immediately neutralized by the addition of a high-pH equilibration buffer (Tris-HCl, pH 8.0-9.0). An additional purification step was performed using a MabSelect™ column (Cytiva). The purity and integrity of the resulting IgG fraction were subsequently confirmed by SDS-PAGE analysis and Coomassie staining.

### Antibody characterization

The isotype of the isolated immunoglobulins was determined using the Pro-Detect™ Rapid Antibody Isotyping Assay Kit - Human (Thermo Fisher Scientific, Waltham, MA, USA). Final protein concentration was determined as described above.

### Real Time Cell Analysis

Dynamic monitoring of cell behavior was performed using the iCELLigence system (ACEA Biosciences; now Agilent Technologies), a label-free platform based on microelectronic impedance sensing. Experiments were conducted in 8-well E-Plates, which is characterised by a specialized bottom surface covered with gold micro-electrode arrays. The measurement principle is based on the interaction between the cells and the electrodes: as cells adhere and spread on the gold surface, they act as insulators, increasing the electrical impedance of the circuit. These changes in impedance were converted by the system into a dimensionless parameter known as the Cell Index (CI), that was normalized (Normalized Cell Index, NCI) for the time of the beginning of the treatment. The NCI is directly proportional to the electrode coverage, reflecting the number, proliferation rate, morphology, and attachment strength of the cells. The iCELLigence RTCA Station was maintained within a humidified incubator at 37°C and 5% CO_2_. Preliminary density scouting determined that a seeding density of 15,000 cells/well was optimal for IMR-90 cells. All treatments were performed in triplicate or quadruplicate once the cells reached the exponential growth phase, with complete medium serving as the negative control. Impedance was recorded every hour prior to treatment, post-treatment every 15 minutes for a total duration of 122 hours. Data were analysed using the RTCA Data Analysis Software (ACEA Biosciences).

### MTT assay

Cell viability and metabolic activity were evaluated using the MTT [3-(4,5-dimethylthiazol-2-yl)-2,5-diphenyltetrazolium bromide] colorimetric assay. IMR-90 cells were seeded in 96-well plates at a density of 8000 cells per well of a 96-well plate. Following an initial stabilization period, cells were then treated with purified MDA5 protein, patient-derived antibodies or a combination of both, each at a final concentration of 1 ìg/ml. On the third day of culture (72 h), the medium was replaced with a solution containing 0.45 mg/mL of MTT in fresh medium, and cells were incubated at 37°C for 2 h. Following incubation, the medium was carefully aspirated, and the resulting crystals were solubilized by adding 100 µL of DMSO per well. The absorbance was measured at 570 nm using a SPECTROstar Omega microplate reader (BMG LABTECH, Ortenberg, Germany), and cell viability was expressed as a percentage relative to the untreated control cells.

### Realtime qPCR

For gene expression analysis, IMR-90 cells were cultured in T75 flasks. The conditions tested were the following: untreated negative control (medium only), MDA5 protein (1 μg/ml), anti-MDA5 antibodies (1 μg/ml) and MDA5/anti-MDA5 complex (both at 1 μg/ml).

Following ca. 5 days (122 h) of incubation, cells were harvested and total RNA was extracted using the GeneJET RNA Purification Kit (Thermo Scientific) according to the manufacturer’s instructions. The concentration of isolated RNA was quantified and subsequently reverse-transcribed into cDNA using the PrimeScript™ II 1st strand cDNA Synthesis Kit (Takara Bio). Quantitative PCR was performed on the CFX Duet Real-Time PCR System (Bio-Rad). For each biological sample, a technical triplicate of the cDNA was loaded onto the plate. A panel of 11 target genes was analyzed, with gene expression levels normalized with the actin gene as reference. Raw cycle threshold (Ct) values were exported. The geometric mean of the reference gene (actin) was used for normalization to calculate the ⊗Ct. Results were then normalized to the untreated control group (ÄÄCt, used for statistical analysis) and the fold change in expression determined using the 2^-ÄÄCt^ method. The primer/probe sets used for gene expression analysis were commercially available TaqMan® Gene Expression Assays ^29^. The assay IDs were as follows: TLR3 Hs01551078_m1, TLR4 Hs00152939_m1, TLR7 Hs01933259_s1, TLR8 Hs00152972_m1, TLR9 Hs00370913_s1, IFN-α Hs00855471_g1, IFN-β Hs01077958_s1, ET-1 Hs00174961_m1, COLIα1 Hs00164004_m1, MMP-1 Hs00899658_m1, and GAPDH Hs99999905_m1. In addition, human β-actin (hActin) primers were used as a reference gene: forward, 5′-CACCATTGGCAATGAGCGGTTC-3′ and reverse, 5′-AGGTCTTTGCGGATGTCCACGT-3′.

### Enzyme-Linked Immunosorbent Assay (ELISA)

To quantify the secretion of specific cytokines and fibrosis markers, cell culture supernatants were harvested 5 d post-treatment and analyzed using SimpleStep ELISA® kits (Abcam) for Interferon-alpha (IFN-α), Interferon-beta (IFN-β), Interferon-gamma (IFN-γ), and Pro-collagen. The assays were performed according to the manufacturer’s instructions using a streamlined immunocapture strategy. Briefly, 50 μl of standards or samples (supernatants from the aforementioned IMR-90 cell pellets) were added to the 96-well microplates, followed by 50 μl of the specific Antibody Cocktail. Each plate included two blank wells (zero control). All cell supernatants were assayed in technical triplicate. After a 1 h incubation at room temperature on a plate shaker (400 rpm), the wells were washed three times with 350 μl of 1X Wash Buffer PT. Subsequently, 100 μl of TMB Development Solution was added and incubated for 10 min in the dark, followed by the addition of 100 μl of Stop Solution. The optical density (OD) was measured at 450 nm using a SPECTROstar Omega microplate reader (BMG LABTECH, Ortenberg, Germany). Data analysis was performed by subtracting the average absorbance of the blank control from all readings.

### Statistical analysis

Statistical data analysis was performed through Graphpad Prism (version 8.0, GraphPad software, San Diego, CA, USA). A data normalization check was performed by Shapiro-Wilk test. ANOVA with post-hoc Tukey test used when negative or Kruskal-Wallis test followed by Dunn’s test when positive were used to assess the statistical differences between samples. A p-value <0.05 was considered statistically significant. Data are reported as the mean±SD or mean±SEM. Every sample was used in triplicate, and each experiment was repeated twice

## 3. Results

### MDA5 Protein Purification and Affinity Resin Preparation

The first phase of the study focused on the production of high-quality recombinant human MDA5 protein. The HEK293F mammalian expression system was strategically selected to ensure that the protein maintained its native conformational integrity and essential post-translational modifications, which are critical for accurate autoantibody recognition. As shown in Fig. 1(a), SDS-PAGE analysis confirmed the success of this strategy.

**Fig. 1.**
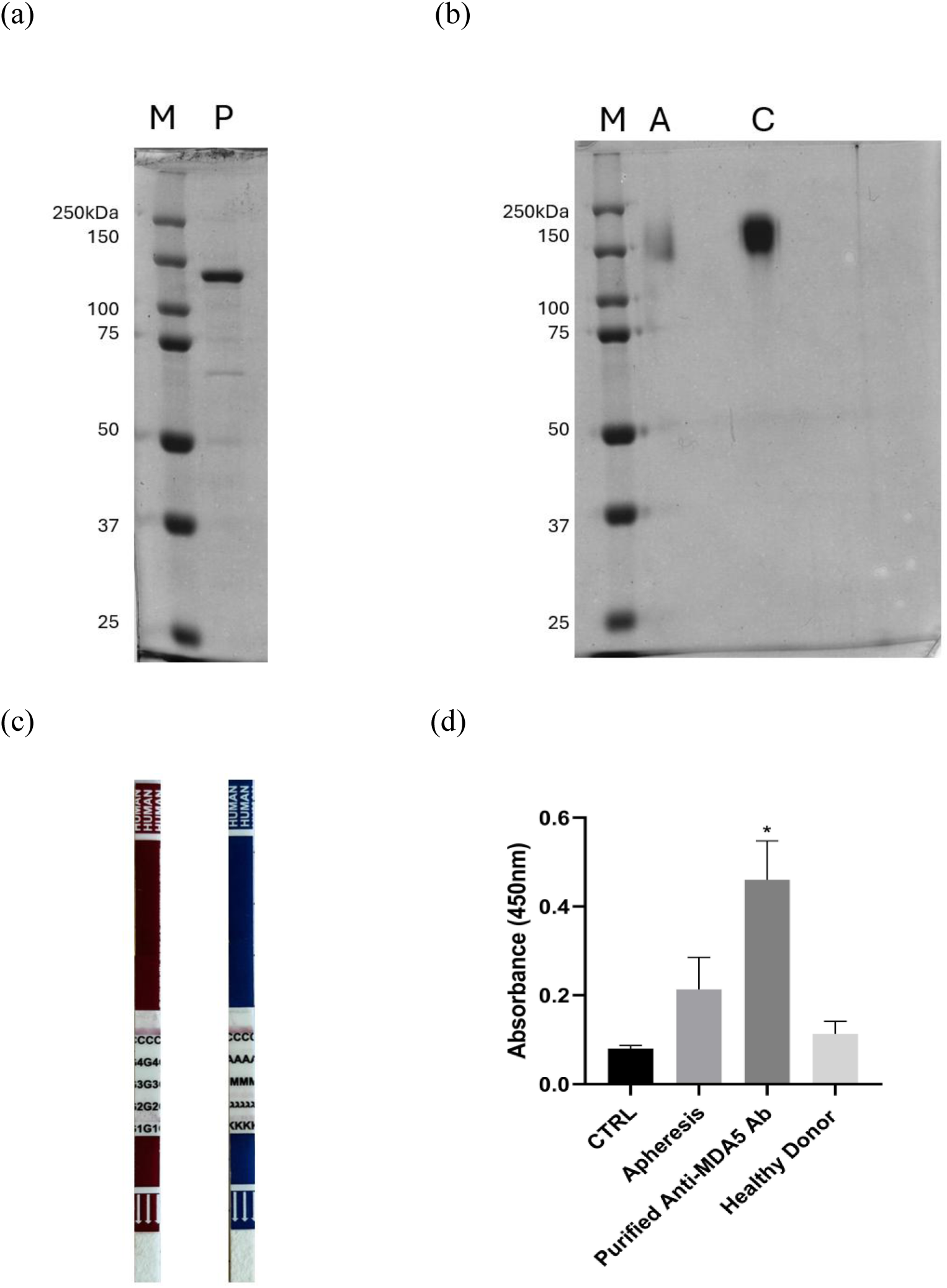
(a) Lane M: Molecular weight maker (Precision Plus Protein standards, Biorad); lane P: purified MDA5. (b) Coomassie gel of anti-MDA5 antibodies: M: Marker; A: Antibodies; P: MDA5 protein; C: Control (Trastuzumab). (c) Antibody isotyping strips (Pro-Detect™ Rapid Antibody Isotyping Assay Kit - Human, Thermo Fisher Scientific, Waltham, MA, USA). Brown: IgG heavy chain identification. Blue: IgM, IgA and light chain identification. (d) ELISA results for MDA5 binding of the original apheresis, purified antibodies and healthy donor serum. *: p=0.014.

### Isolation of polyclonal antibodies from patient’s serum

Therapeutic plasmapheresis was performed as part of the clinical management of the patient. Autoantibodies were selectively isolated using the antigen-specific approach described above. This strategy allowed the enrichment and purification of anti-MDA5 autoantibodies with high specificity (Fig. 1d), enabling downstream analyses aimed at investigating their functional and pathogenic properties. The isotype turned out to be mainly IgG1, with both κ and λ light chain identity (Fig. 1c). The mean antibody recovery was 415±43 µg per 50 ml of apheresis.

### Real Time Cell Analysis

All curves of the RTCA progressively increased, which was consistent with continued proliferation and/or spreading of IMR90 fibroblasts (Fig. 2a) and suggested that all treatments were generally well tolerated by IMR90 cells. In fact, they mainly affected the rate of increase of the Normalized Cell Index (NCI) after the beginning of the treatment (vertical black line, t=24 h), where the NCI is an index of the proliferative/adhesive behavior of cells. The untreated control (red line) showed the lowest NCI at later time points. The MDA5-treated cells (green line) and the anti-MDA5 antibody condition (blue line) displayed an NCI only moderately higher than the control from approximately 60–70 h onward. This suggested that both MDA5 and the anti-MDA5 antibodies had a limited effect on cells. Interestingly, the combination of MDA5/anti-MDA5 antibodies (pink line) showed the highest NCI throughout the later phase of the experiment and increasingly diverged from the control over time, reaching an approximately 25% increase compared to control at t=121 h (p=0.0182, Fig. 3a). Hence, the presence of antibodies did not appear to neutralize the MDA5 effect, but to determine a further increase in the RTCA signal compared to the antibodies and MDA5 alone, indicating an effect on proliferation, cell spreading/adhesion, or in inducing a more activated fibroblast phenotype. This might indicate an additive or synergistic effect between MDA5 and anti-MDA5 antibodies. The cell doubling time (Fig. 3b) was inversely related to the NCI at 121 h (Fig. 3a), with a reduction by about 8 h in the MDA5/anti-MDA5 sample value with respect to the control (54.41±0.52 h versus 46.76±0.33 h, p=0.018).

**Fig. 2.**
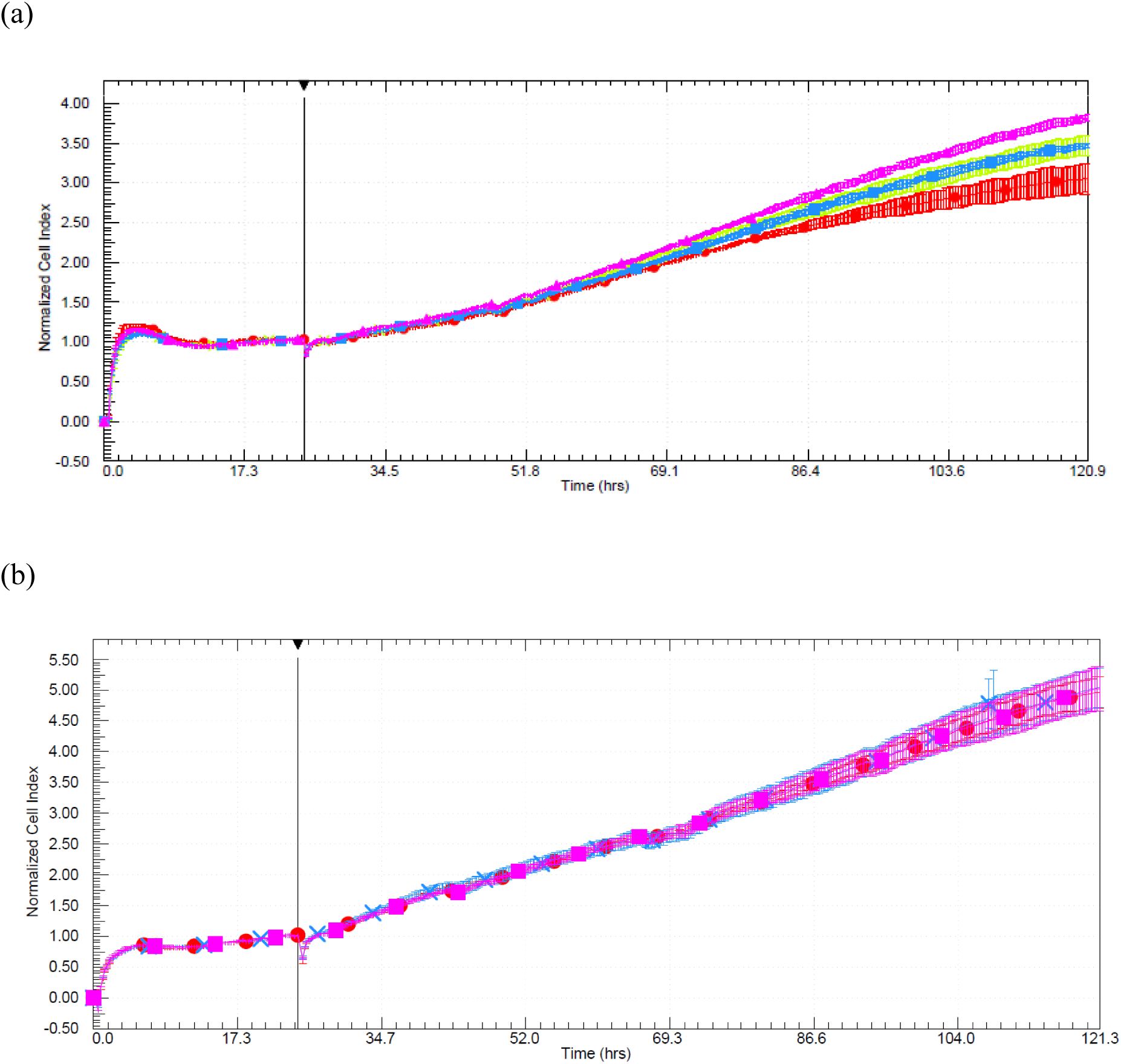
RTCA results: (a) Normal serum; (b) Inactivated serum. Legend: red: control; blue; anti-MDA5 antibodies; green: MDA5; magenta: MDA5/anti-MDA5 antibodies. Lines represent the average ± SD of at least two independent wells.

**Fig. 3.**
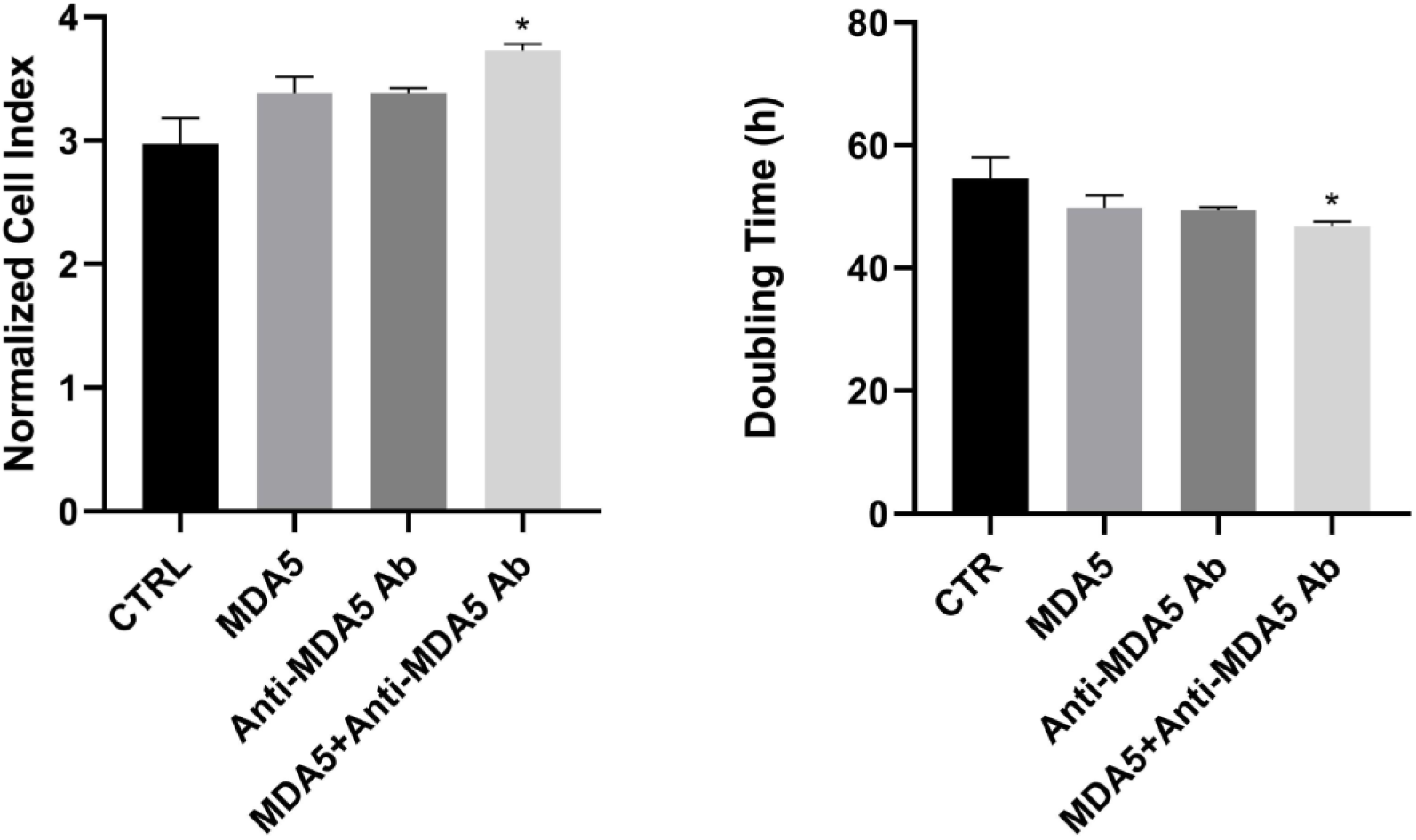
(a) NCI at 121 h (normal serum). (b) Cell doubling time (normal serum). *p<0.05.

Removal of complement (Fig. 2b) completely abolished the effect of the treatment with anti-MDA5 antibodies, both in the presence and in the absence of MDA5.

### Evaluation of Metabolic Activity and Cell Proliferation

Cellular metabolic viability was quantitatively assessed using the MTT assay after 72 h of exposure to the various experimental conditions. As illustrated in Fig. 4a, all tested conditions maintained high levels of viability and statistical analysis confirmed the absence of a significant cytotoxicity across all treatment groups. Specifically, the comparison between the untreated control (CTR) and the cells exposed to purified MDA5 protein alone showed no relevant variation (p=0.935), indicating that the antigen itself did not alter cellular survival. The group treated with patient-derived anti-MDA5 antibodies showed a slight downward trend in viability; however, this did not reach a full statistical significance (p=0.060 vs. CTR). Similarly, the combination of MDA5 and anti-MDA5 antibodies (immune complexes) resulted in viability levels entirely comparable to the control (p=0.5238).

**Fig. 4.**
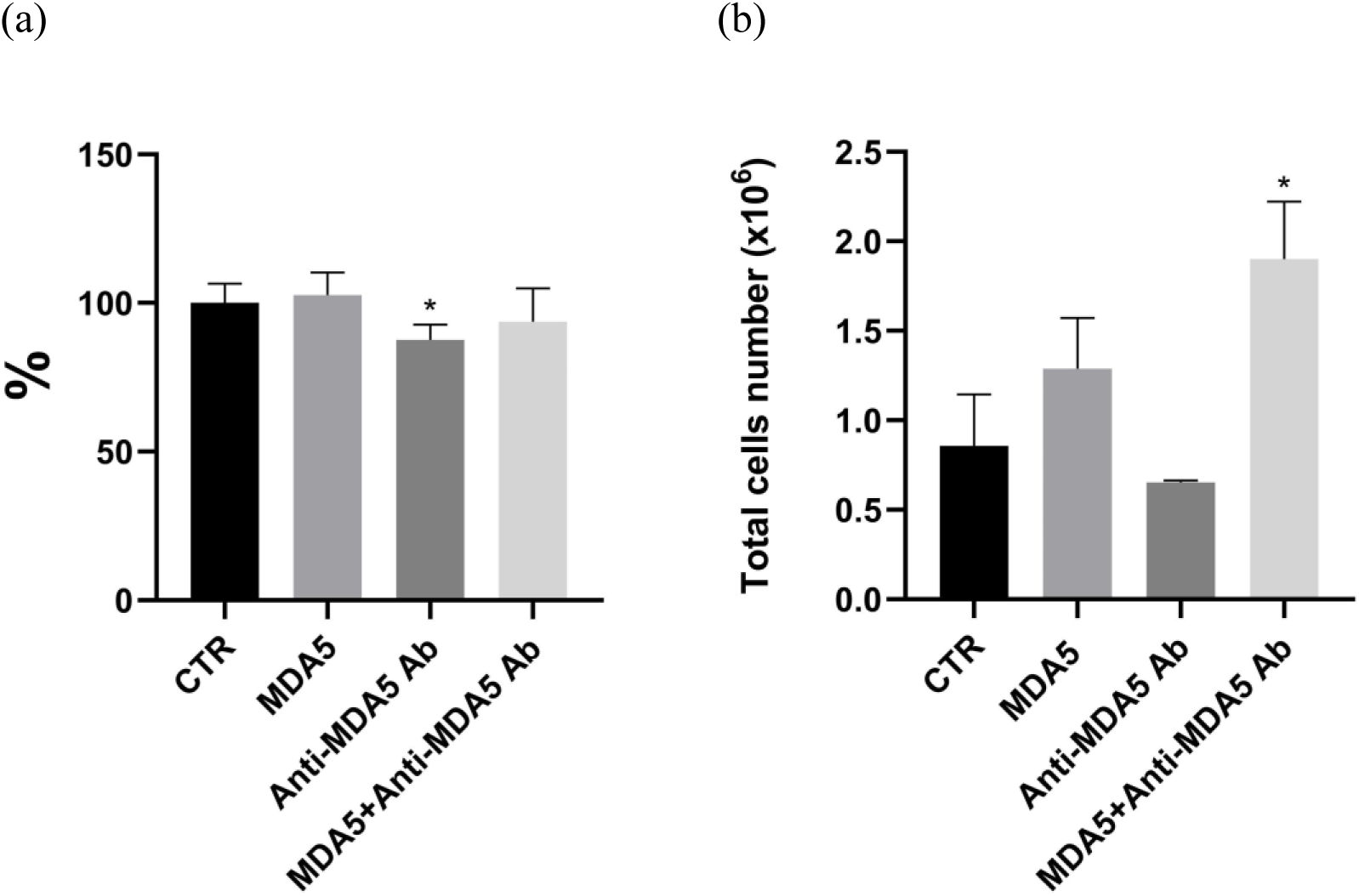
(a) MTT assay. Cells were treated with purified MDA5, patient-derived anti-MDA5 antibodies (Ab), or their combination for 72 h. Data are expressed as a percentage of viability relative to the untreated control (mean ± SD, *: p=0.0606). (b) Total cell number in flask (mean ± SD) at 5 d of treatment (*: p=0.0123).

**Fig. 5.**
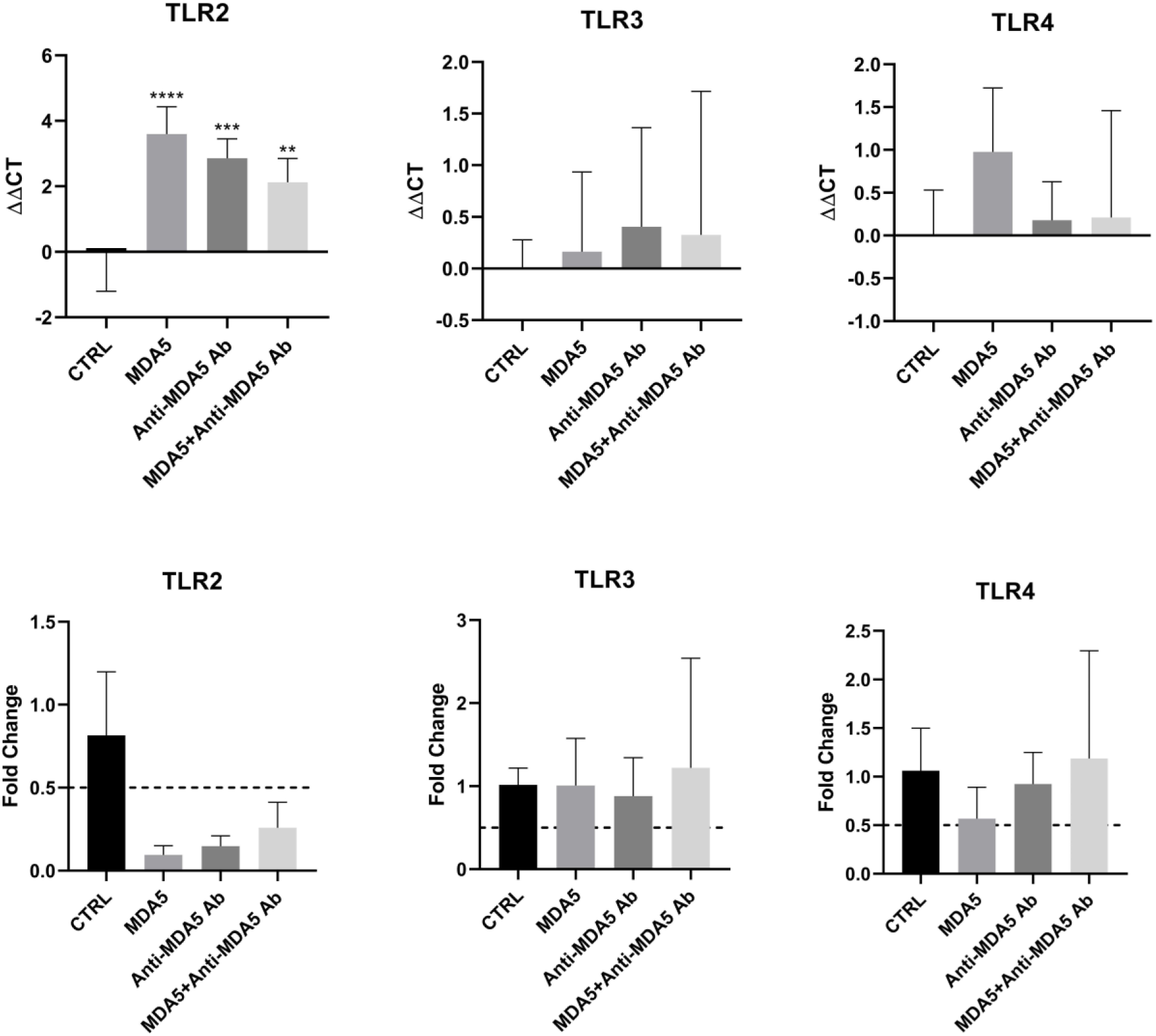
*tlr2, tlr3* and *tlr4* mRNA expression levels in IMR90 lung fibroblasts stimulated with MDA5, anti-MDA5 antibodies or a mixture of the two, at a concentration of 1 μg/ml each. **: p < 0.001, ***: p < 0.0001 versus untreated control.

**Fig. 6.**
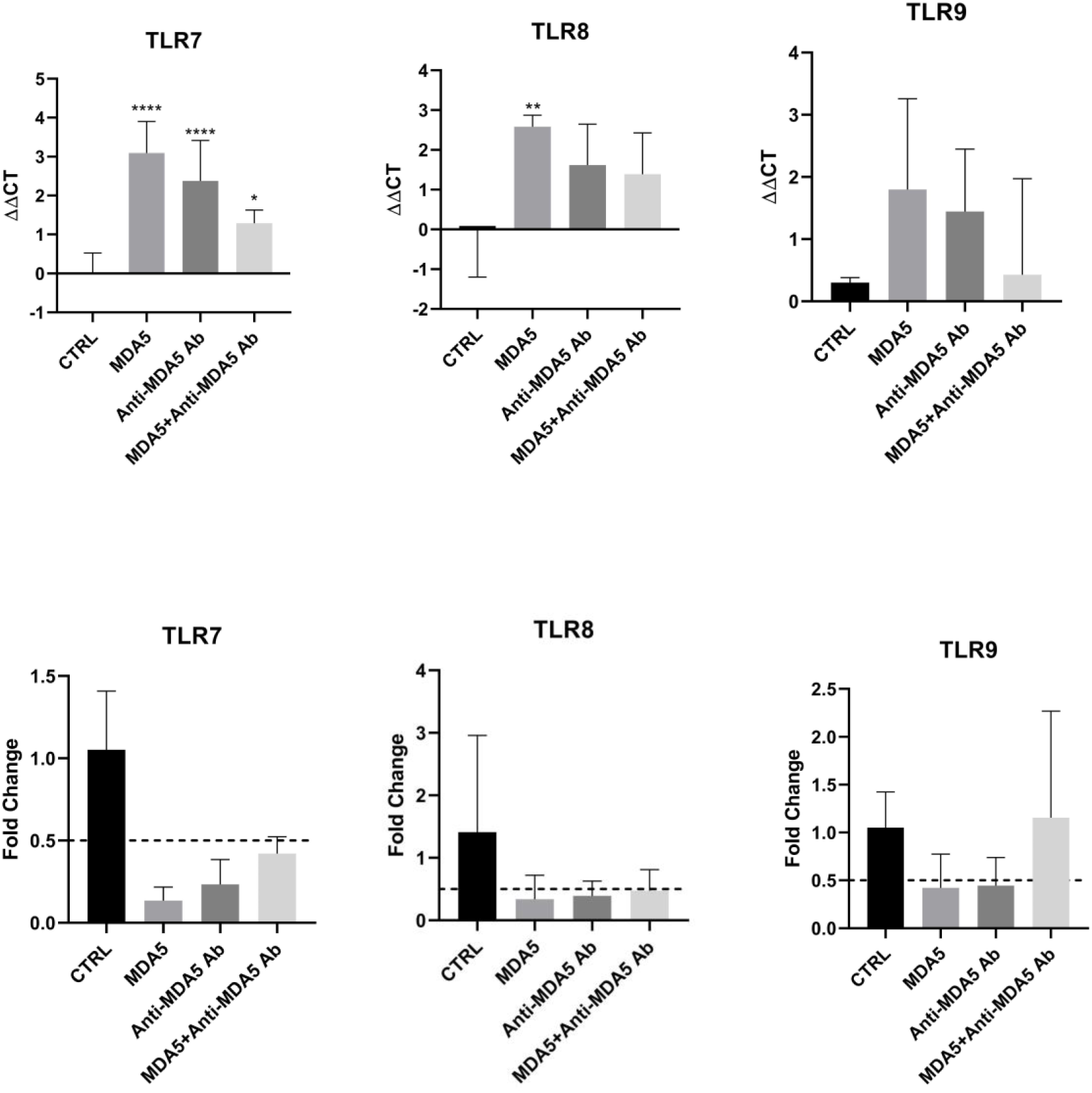
*tlr7, tlr8* and *tlr9* mRNA expression levels in IMR90 lung fibroblasts stimulated with MDA5, anti-MDA5 antibodies or a mixture of the two, at a concentration of 1 μg/ml each. *: p < 0.01, **: p < 0.001, ***: p < 0.0001 versus untreated control.

**Fig. 7.**
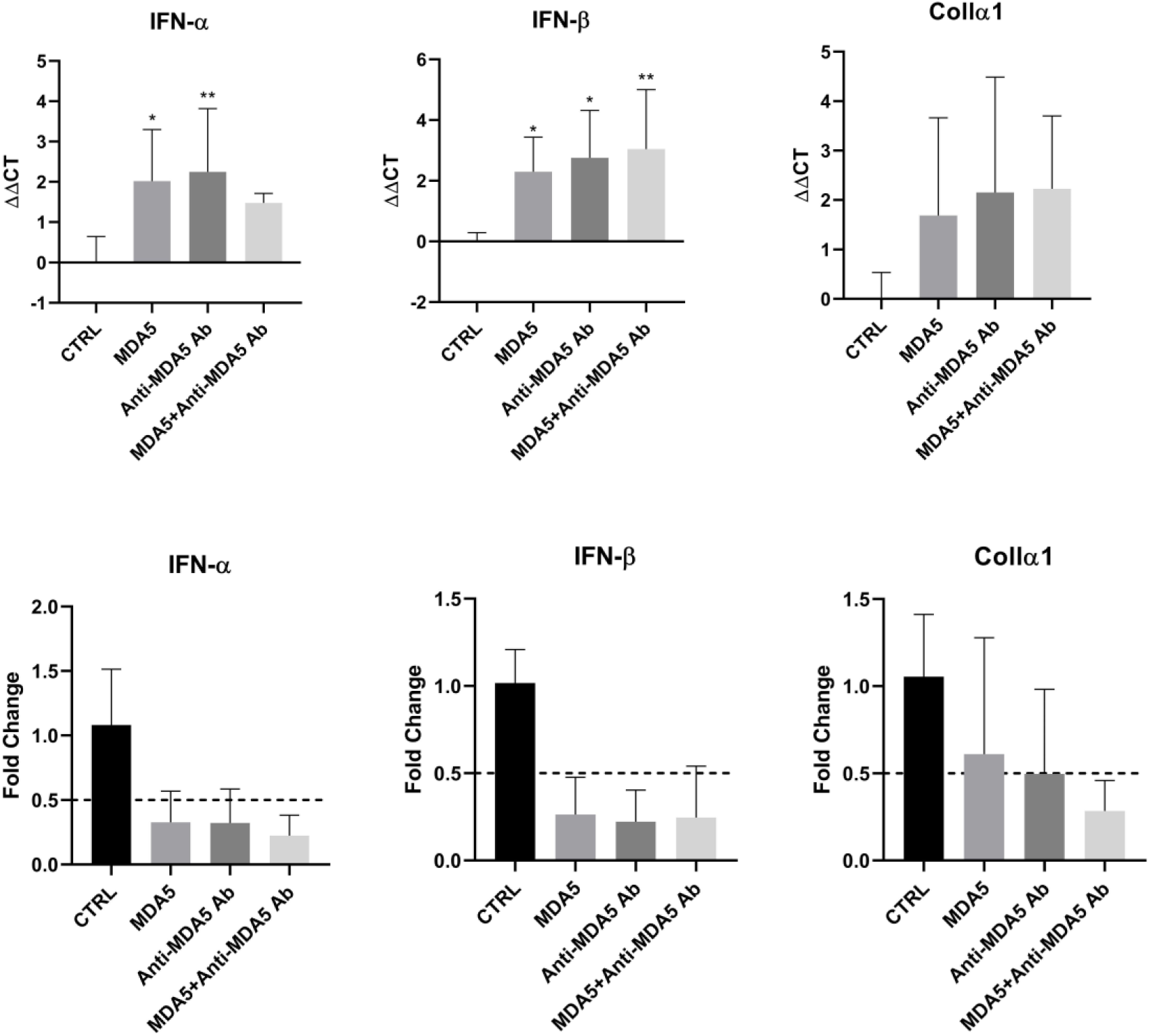
*ifn-α, ifn-β* and *collα1* mRNA expression levels in IMR90 lung fibroblasts stimulated with MDA5, anti-MDA5 antibodies or a mixture of the two, at a concentration of 1 μg/ml each. *: p < 0.01, **: p < 0.001 versus untreated control.

**Fig. 8.**
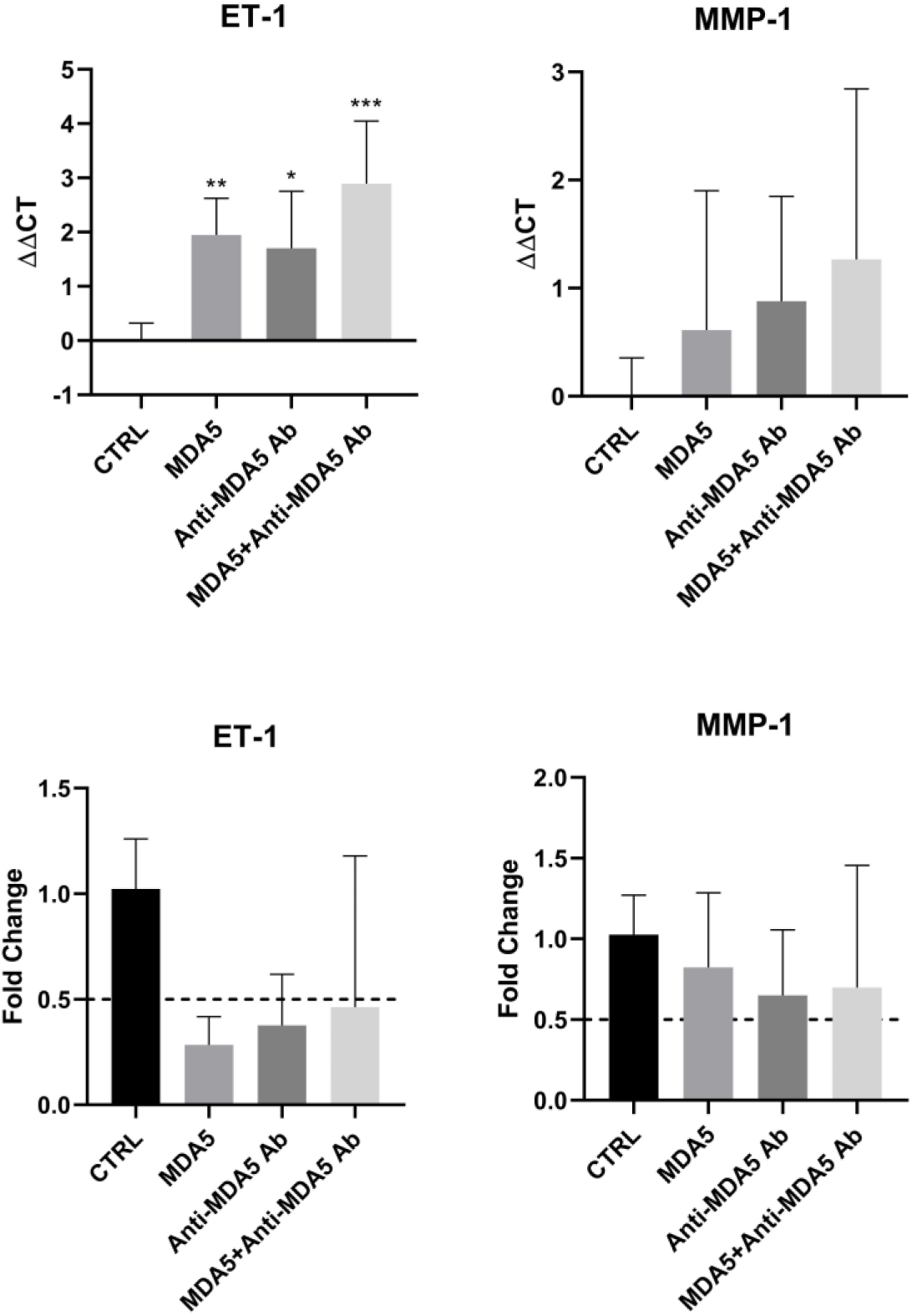
*et-1* and *mmp-1* mRNA expression levels in IMR90 lung fibroblasts stimulated with MDA5, anti-MDA5 antibodies or a mixture of the two, at a concentration of 1 μg/ml each. *: p < 0.01, **: p < 0.001, ***: p < 0.0001 versus untreated control.

**Fig. 9.**
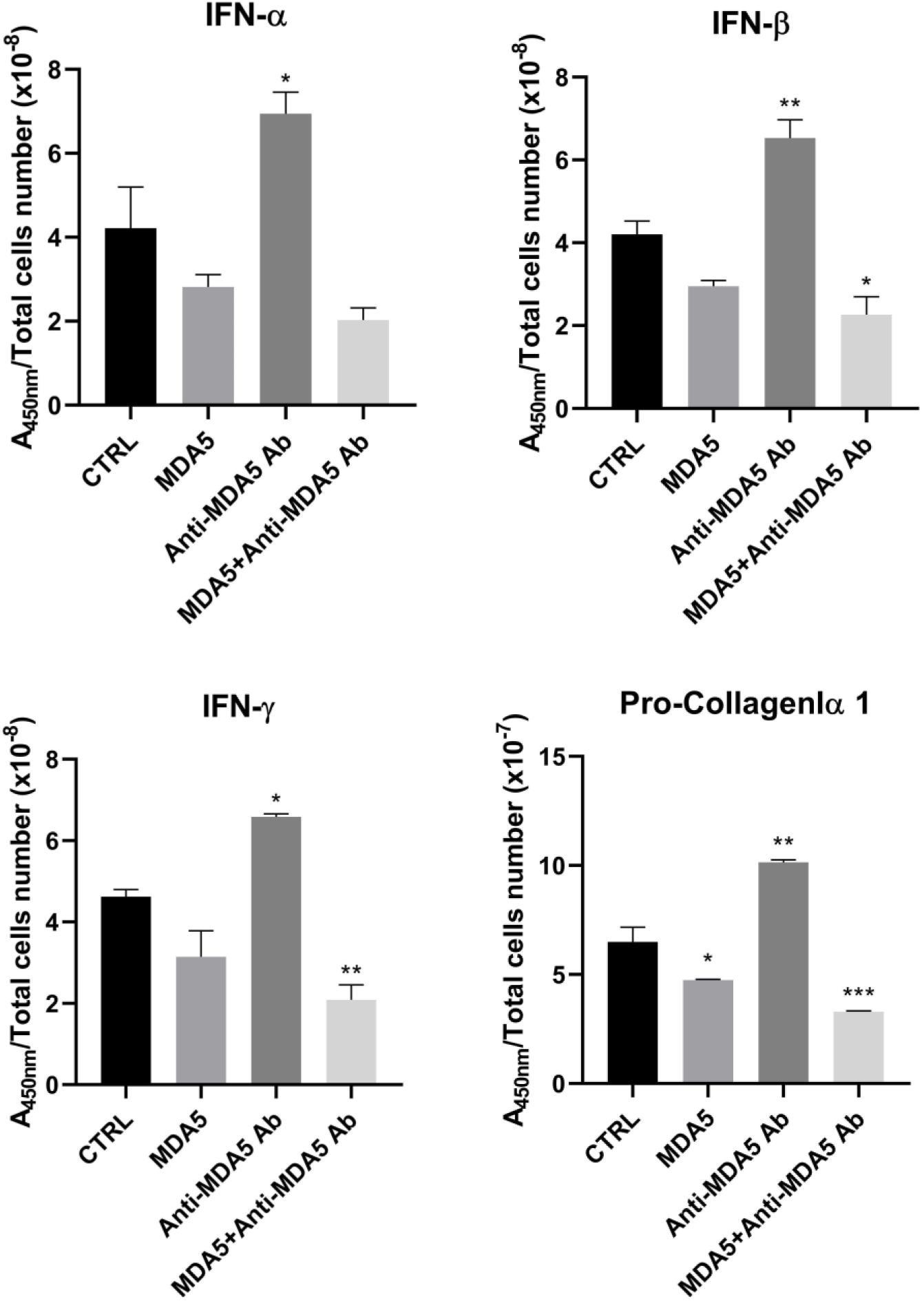
IFN-ɑ, IFN-β, IFN-ɣ and pro-collagen ɑ1 secretion in IMR90 fibroblasts stimulated with MDA5, anti-MDA5 antibodies or a mixture of the two, at a concentration of 1 μg/ml each. *: p < 0.01, **: p < 0.001, ***: p < 0.0001 versus untreated control.

An experiment similar to the one performed for RTCA was scaled up by culturing cells in T75 flasks and counting them at 5 d after treatment (Fig. 4b). The results showed a statistically significant increase (p=0.0123) in cell number in the sample treated with the MDA5/Anti-MDA5 Ab combination, with a number of cells approximately doubled compared to the control, thus supporting a pro-proliferative effect consistent with the RTCA results.

### Real time qPCR

Molecular analysis performed via RT-qPCR after 5 d of incubation revealed significant alterations in the gene expression profile of IMR-90 fibroblasts across various experimental conditions. Specifically, a statistically significant downregulation (p<0.05) was observed in all treated groups compared to the untreated control group for TLR2, TLR7, IFN-β and endothelin-1 (ET-1). No significant differences were observed for TLR3, TLR4, TLR9, collagen ɑ1 and MMP1 (p>0.05), while, for TLR8, only MDA5 seems to determine a significant reduction in mRNA transcription versus the control. MDA5 and anti-MDA5 antibodies alone were also able to induce a reduction in expression of IFN-ɑ. The reliability of these findings is underscored by the normalization of data against the internal reference gene, actin, which ensured consistency across independent experimental sets and confirmed that the treatments drive a specific pro-pathogenic reprogramming of the fibroblasts.

### ELISA

Complementing the transcriptomic data, ELISA assays on day 5 supernatants were utilized to quantify the secretion of specific cytokines and fibrosis markers in the culture supernatants. Data were normalized for the number of cells counted on day 5. Following the treatments, anti-MDA5 antibodies were the most active in inducing a statistically significant increase in IFN-ɑ, IFN-β, IFN-ɣ and pro-collagen ɑ1, while samples exposed to the MDA5/anti-MDA5 immunocomplexes showed a reduction in the levels detected in the supernatant. The increased production of pro-collagen and its modulation by the presence of MDA5 is of particular clinical importance, as it provides direct evidence of active extracellular matrix synthesis, a hallmark of the lung tissue remodeling observed in rapidly progressive interstitial lung disease.

## 4. Discussion

The clinical management of anti-MDA5 syndrome is heavily influenced by its frequent and severe association with rapidly progressive interstitial lung disease (RP-ILD) ^12,13^. While anti-MDA5 autoantibodies have long been utilized as essential diagnostic and prognostic biomarkers ^14,15^, their potential role as active participants in the underlying pathophysiology of lung injury has remained largely speculative. The results of the present study provide mechanistic evidence that these autoantibodies act as direct potential pathogenic drivers that stimulate human lung fibroblasts toward a proliferative pro-fibrotic phenotype. The shift of resident fibroblasts from a quiescent state to an activated, contractile myofibroblast phenotype is a central event in fibrotic lung remodeling, where persistent myofibroblast activation drives excessive extracellular matrix deposition and irreversible architectural distortion ^16,17^. In this view, the selection of the IMR-90 fetal lung fibroblast cell line provided a robust and sensitive model to investigate these interactions, as these cells maintain a responsive phenotype capable of reacting to inflammatory and fibrotic stimuli ^18,19^. Our initial functional assessment using RTCA demonstrated that MDA5, the patient’s anti-MDA5 antibodies and the interaction between them significantly altered fibroblast behavior. The observed increase in the Normalized Cell Index (NCI) and the corresponding reduction in doubling time, already present with each of the single treatments, was most pronounced when cells were exposed to the MDA5/anti-MDA5 immune complexes, suggesting a summation effect that excludes neutralization of the antigen action by the antibodies. This suggests that the antibody itself influences IMR90 behavior, possibly through Fc-mediated effects, receptor interactions, immune-like signaling pathways or modulation of endogenous MDA5-related signaling. In this respect, our observation that serum without complement (complement-inactivated) completely abolished the fibroblast RTCA response to anti-MDA5 antibodies with or without MDA5, underscores the role of the Fc in activation of the classical complement pathway as an essential mediator. This suggests that immune-complex and complement deposition in the lungs truly contributes to local tissue injury in anti-MDA5–positive DM-ILD and does not represent only an epiphenomenon, like already demonstrated in other autoimmune conditions ^20^⁻^22^. The presence of C3 in ILD patients lung tissue is consistent with this in vitro observation ^20^. There is, in fact, strong evidence for local complement activation in the lung, but relatively limited and inconsistent evidence regarding systemic complement consumption in anti-MDA5 dermatomyositis.

Because RTCA measures electrical impedance, an increased NCI does not in general exclusively mean an increased cell number. This impedance-based signal reflects a combination of increased cell proliferation, enhanced substrate adhesion, and morphological spreading, which are characteristic of fibroblast activation in pulmonary remodeling ^23^⁻^25^. Crucially, the MTT assays confirmed that these changes were not a consequence of acute cellular stress or toxicity. All treatment conditions maintained high levels of metabolic viability, indicating that the observed molecular changes represent a specific functional reprogramming rather than a non-specific response to injury. The statistically significant increase in cell numbers following treatment with the immunocomplex, associated to the reduction of the doubling time, further reinforces the hypothesis that these antibodies promote a fibroproliferative environment. This is particularly relevant in the clinical context of RP-ILD, where the rapid accumulation of activated fibroblasts and the subsequent deposition of extracellular matrix lead to a swift decline in pulmonary function. The molecular basis for this activation was further elucidated through RT-qPCR and ELISA analyses, which revealed a significant alteration in the levels of key genes and proteins involved in both innate immunity and fibrotic progression.

An unexpected finding of our study was the significant reduction in the mRNA levels of several innate immune and inflammatory genes, including TLR2, TLR7, IFN-α, IFN-β and ET-1, despite the concurrent increase in fibroblast proliferation, RTCA signal, IFNs and pro-collagen production. This apparent discrepancy may reflect the transition from an early inflammatory response to a fibroproliferative state, a phenomenon increasingly recognized in chronic fibrotic disorders such as idiopathic pulmonary fibrosis and systemic sclerosis. Activated fibroblasts are known to acquire a phenotype characterized by enhanced proliferation, extracellular matrix production, and resistance to apoptosis, while simultaneously exhibiting reduced responsiveness to inflammatory stimuli ^16,17^. Furthermore, prolonged stimulation of RNA-sensing pathways may induce compensatory negative-feedback mechanisms leading to downregulation of Toll-like receptors and interferon-related genes, a process described for both TLR and RIG-I-like receptor signaling pathways. Since gene expression was assessed after five days of continuous stimulation, it is possible that an initial inflammatory burst occurred at earlier time points and was subsequently followed by receptor desensitization and transcriptional repression, whereas secreted proteins accumulated in the culture supernatant over time. The use of IMR-90 fetal lung fibroblasts may also have contributed to this response profile. Compared with adult fibroblasts, fetal fibroblasts display greater proliferative capacity, enhanced tissue-remodeling properties, and distinct innate immune responses, favoring repair and developmental programs over sustained inflammation. Therefore, the observed reduction in inflammatory transcripts may not indicate an absence of pathogenic activation, but, rather, a shift toward a profibrotic, tissue-remodeling phenotype that may be relevant to the pathogenesis of anti-MDA5-associated interstitial lung disease. In this respect, the detection of increased pro-collagen levels induced by anti-MDA5 antibodies is perhaps the most direct evidence of the pro-fibrotic potential. Pro-collagen secretion is a definitive marker of active extracellular matrix synthesis, which translates to the loss of gas exchange capacity in a clinical setting. The fact that these responses were observed using patient-derived antibodies lends significant weight to the argument that therapeutic strategies targeting these circulating factors, such as plasmapheresis, are mechanistically sound.

Taken together, the RT-qPCR and ELISA results demonstrate that anti-MDA5 autoantibodies are not merely diagnostic markers but act as direct pathogenic drivers, a finding of the present study that aligns with a growing body of evidence in other autoimmune diseases. Research in systemic sclerosis (SSc) has demonstrated that immune complexes containing SSc-specific autoantibodies—such as those against topoisomerase I or RNA polymerase III—induce a distinct proinflammatory and profibrotic phenotype in skin fibroblasts ^29^. SSc-ICs were found to alter innate immune sensors, significantly upregulating the expression of Toll-like receptors (TLRs) and stimulating the secretion of pro-collagen and endothelin-1 (ET-1) in patient derived fibroblasts after 24-48 h incubation^29^. This direct pathogenic role for circulating immune complexes provides a robust mechanistic rationale for the clinical efficacy of plasmapheresis, which is utilized to treat both severe SSc and rapidly progressive anti-MDA5-associated ILD by removing these pathogenic factors from the circulation.

The isotype characterization of the anti-MDA5 autoantibodies isolated in our study revealed a predominant IgG1 subclass, which aligns with established clinical data identifying IgG1 as the most prevalent subclass in the anti-MDA5 dermatomyositis subset ^30,31^ . This finding is of particular pathogenic relevance, as the IgG1 isotype is a potent activator of the classical complement pathway ^32^. This supports our observation that complement-inactivated serum completely abolished the activation of IMR-90 fibroblasts in the RTCA model, reinforcing the hypothesis that the pathogenic effects of these autoantibodies are mediated through the formation of complement-fixing immune complexes. Furthermore, our detection of both kappa and lambda light chains reflects the polyclonal nature of the autoantibody response in these patients. This is consistent with recent single-cell repertoire analyses of MDA5-reactive B cells, which have identified multiple distinct B cell lineages ^33,34^.

A crucial prerequisite for the formation of the pathogenic immune complexes (ICs) described in our study is the extracellular availability of the MDA5 antigen, a protein traditionally regarded as strictly cytosolic. Recent evidence has confirmed the existence of a soluble form of the MDA5 protein in human sera, detectable in both healthy individuals and patients with dermatomyositis ^35^. Notably, this soluble MDA5 is not merely a consequence of passive leakage from necrotic cells; research indicates it can be rapidly released from the cytoplasm of peripheral blood mononuclear cells within 15 min of double-stranded RNA stimulation, appearing as both a full-length 120 kDa protein and a 60 kDa fragment containing the helicase domain ^35^. This extracellular presence provides the biochemical basis for the formation of circulating MDA5/anti-MDA5 immunocomplexes. Furthermore, the strong expression of MDA5 in alveolar macrophages in normal lung ^35^ and its robust expression in the alveolar epithelium and macrophages of DM-ILD lungs and MDA5-transgenic injury models ^14^, together with the demonstrable release of soluble MDA5 from stimulated immune cells ^35^, support the notion that the pulmonary microenvironment is a major source and potential reservoir of soluble MDA5 antigen in anti-MDA5 dermatomyositis. Furthermore, a unique subset of CD14^+^ monocytes that highly expressed MDA5 was specifically identified in anti-MDA5 patients ^33^.

## 5. Conclusions

To our knowledge, this is the first time human anti-MDA5 specific antibodies have been purified, characterized and used to test their effect on cultured fibroblasts. Though the anti-MDA5 antibodies we used derive from a single patient, all our findings are consistent with other observations reported in the literature and suggest that the pathogenesis of RP-ILD in anti-MDA5-positive patients is driven by a complex interplay where autoantibodies trigger a specific, complement-mediated, pro-pathogenic program in lung fibroblasts, contributing to the progressive remodeling characteristic of this severe dermatomyositis subset. This study also represents the first experimental attempt to evaluate the direct biological effects of purified, recombinant MDA5 protein on a cellular model. While MDA5 is traditionally characterized as an intracellular sensor, our approach, testing the purified molecule both in isolation and within immune complexes, provides a novel mechanistic framework, demonstrating how the extracellular presence of the MDA5 molecule itself can act as a primary trigger for the pro-pathogenic reprogramming of lung fibroblasts.

## Funding

This work was supported by the 2022 Progetti di Rilevante Interesse Nazionale (PRIN) programme (code 20224H9JW9).

## Acknowledgements

We thank Dr. Cesare Perotti for collaboration with the preparation of the plasmapheresis and Prof. Sun Hur for kindly providing the human MDA5 cDNA.

## Notes

### Competing Interest Statement

The authors have declared no competing interest.

